# Quantum transport in mitochondrial complex I is governed by a conserved structural bottleneck

**DOI:** 10.64898/2026.05.28.728423

**Authors:** Ji-Yong Sung, Jae-Ho Cheong

**Affiliations:** Department of Neurosurgery, Seoul National University Bundang Hospital, Seoul National University College of Medicine, Seongnam-si, Republic of Korea; Department of Surgery, Yonsei University College of Medicine, Seoul 03722, Republic of Korea; Department of Quantum Information Science, Graduate School, Yonsei University, Seoul 03722, Republic of Korea

## Abstract

Electron transport in mitochondrial complex I is mediated by a chain of redox centers, yet how electrons traverse this network beyond the canonical pathway remains unclear. While prior models treat transport as a sequential process, they do not resolve whether alternative pathways contribute to functional electron flow. Here, we formulate electron transport as a continuous-time quantum walk on a structure-derived redox network and systematically map pathway-level electron flux inferred from quantum-walk dynamics across species. We identify a conserved structural bottleneck at the N5–N6a interface that suppresses direct electron transfer. Strikingly, quantum-walk flux analysis indicates that this bottleneck does not simply limit transport, but can redistribute electron flux into residue-mediated alternative pathways. Across species, these alternative routes support substantial flux and, in several cases, are comparable to or can exceed the canonical direct pathway, indicating a conserved mechanism of pathway-level flux redistribution. This behavior arises from geometric constraints encoded in protein structure and persists under environmental decoherence, demonstrating that architecture governs not only transport efficiency but also the organization of electron flow within the network. Together, our findings suggest a network-level organization of electron transport in complex I, in which a structurally encoded bottleneck reshapes flux through alternative pathways, consistent with a structurally encoded link between protein geometry and quantum transport behavior. We note that the bottleneck-dominated and flux-redistribution observations are not in tension: suppression of the direct N5–N6a step is precisely what redirects amplitude into the parallel residue-mediated routes.

## Introduction

Electron transport is a fundamental process that underlies cellular energy conversion, enabling the coupling of redox reactions to ATP synthesis across all domains of life.^1,2^ In mitochondria, complex I (NADH:ubiquinone oxidoreductase) functions as the primary entry point of electrons into the respiratory chain, catalyzing electron transfer from NADH to ubiquinone through a chain of flavin and iron–sulfur (Fe–S) redox centers. This process spans distances of up to several nanometers and involves multiple sequential electron-transfer steps embedded within a large protein complex exceeding 40 subunits in eukaryotes.^3-5^ Despite decades of structural, biochemical, and computational studies, the principles that determine the efficiency of electron transport in complex I remain incompletely understood. ^6^ While high-resolution cryo-electron microscopy has revealed the architecture of the redox chain in unprecedented detail, linking this structural information to functional transport efficiency remains a central challenge in bioenergetics. At the level of individual electron-transfer events, rates are often described by Marcus theory, which relates transfer kinetics to electronic coupling, reorganization energy, and thermodynamic driving force.^7,8^ However, in extended multi-step systems such as complex I, electron transport is not governed by a single transfer event but emerges from a sequence of coupled hopping processes.^4,9-11^ In such systems, it is unclear whether transport efficiency is distributed across many steps or dominated by specific rate-limiting transitions.

In addition to the question of rate limitation, an equally fundamental issue remains unresolved: whether electron transport in multi-step redox systems is confined to a single canonical pathway or can be supported by alternative routes within the network. In complex I, the redox centers are embedded in a three-dimensional protein architecture, where multiple residue-mediated connections could, in principle, provide additional channels for electron transfer. However, conventional sequential and hopping-based models inherently assume a dominant linear pathway and do not capture how network topology may enable alternative flux propagation. As a result, it remains unclear whether such alternative pathways can contribute meaningfully to electron transport at the network level or how their contribution is shaped by underlying structural constraints.

This raises a fundamental question: What determines the overall efficiency of multi-step electron transport in biological systems? In particular, it remains unresolved whether transport efficiency is primarily dictated by local properties, such as the chemical environment, polarity, or protein dynamics surrounding individual redox centers, or by global structural constraints encoded in the spatial organization of the redox chain.^12^ Resolving this question is essential for understanding not only mitochondrial bioenergetics but also general principles of long-range electron transfer in biological and synthetic systems. ^4,10,11 13^

Recent advances in structural biology have made it possible to compare complex I architectures across diverse species, revealing a broadly conserved arrangement of redox centers alongside subtle variations in inter-cluster distances and local environments. At the same time, theoretical and experimental studies have suggested that quantum effects, including coherence and environmental decoherence, may influence transport efficiency in biological electron transfer systems. In particular, the concept of environment-assisted quantum transport has emerged as a potential mechanism by which biological systems optimize energy transfer under noisy conditions.^14,15^ However, whether such effects operate in large-scale redox chains such as complex I, and how they relate to structural constraints, remains largely unexplored. ^16^

Notably, atomistic studies have already mapped the inter-cluster tunneling pathways of complex I in detail, identifying the N5–N6a step as the longest, rate-limiting transfer and resolving the contributing cysteine-mediated bridges at single-residue resolution. Building on this structural foundation, our aim here is distinct and complementary: rather than computing single-step couplings in one organism, we ask a network-level, cross-species question— whether system-level transport collapses onto a single structurally encoded constraint, and how that constraint reorganizes flux across the redox network. Here, we address these questions by systematically analyzing electron transport in mitochondrial complex I across ten species spanning bacteria to mammals within a continuous-time quantum-walk framework.^17^ Using structure-derived electronic couplings, we show that electron transport is universally governed by a single dominant kinetic bottleneck along the redox chain.^10^ We further demonstrate that this bottleneck is structurally encoded, arising from inter-cluster distance rather than local chemical environment, and quantitatively determines transport efficiency across species. ^9 18^ By incorporating environmental decoherence within an open quantum system framework, we further show that transport efficiency exhibits a non-monotonic dependence on environment, consistent with environment-assisted quantum transport.^14^ In addition, quantum-walk analysis reveals that the structural bottleneck is associated with redistribution of electron flux into alternative pathways within the redox network.^10 19^ Together, our findings point to a link between protein architecture, quantum dynamics, and network-level electron transport, in which structurally encoded constraints appear to shape both the efficiency and the organization of electron flow in this system.

## Results

### A structurally conserved kinetic bottleneck governs electron transport in complex I

To determine how electron transport efficiency is distributed along the redox chain of mitochondrial complex I, we quantified step-specific electronic coupling strengths across ten species spanning bacteria to mammals. The electron-transfer pathway was defined as a sequential chain from FMN through a series of Fe–S clusters to the terminal N2 site, and all couplings were normalized to enable direct comparison across species. Across all species examined, we observed a strikingly non-uniform distribution of electronic coupling strengths along the pathway (**Fig. 1**). Rather than being evenly distributed, coupling values exhibit a pronounced minimum at the N5→N6a transition. This minimum is consistently localized to the same structural region across all species analyzed, indicating the presence of a conserved kinetic bottleneck embedded within the architecture of complex I. Importantly, the magnitude of this bottleneck is substantial. The weakest coupling at the N5→N6a step is reduced relative to neighboring steps, generating a sharp kinetic contrast along the pathway. This contrast effectively partitions the redox chain into upstream and downstream segments with distinct transport characteristics, suggesting that electron transfer is funneled through a single dominant constraint rather than distributed across multiple comparable steps. Although the spatial position of the bottleneck is highly conserved, its strength varies across species. To quantify this variation, we expressed coupling values as −log10(J_norm_), where larger values correspond to weaker electronic coupling and stronger kinetic limitation. The resulting heatmap reveals a conserved bottleneck signature centered on the N5→N6a region, with species-specific differences primarily reflected in the magnitude of coupling rather than its location (**Fig. 1**). This pattern indicates that while the structural position of the rate-limiting step is evolutionarily constrained, the degree of transport limitation can be tuned across species. Such variation suggests that evolutionary adaptation operates not by relocating the bottleneck, but by modulating its strength within a fixed structural framework. Because our dataset comprises static structural snapshots of extant species that are not phylogenetically independent, we interpret the conservation of the bottleneck location as an observation rather than direct evidence of adaptive selection; formal tests of adaptation would require phylogenetically corrected comparative analysis.

**Figure 1.**
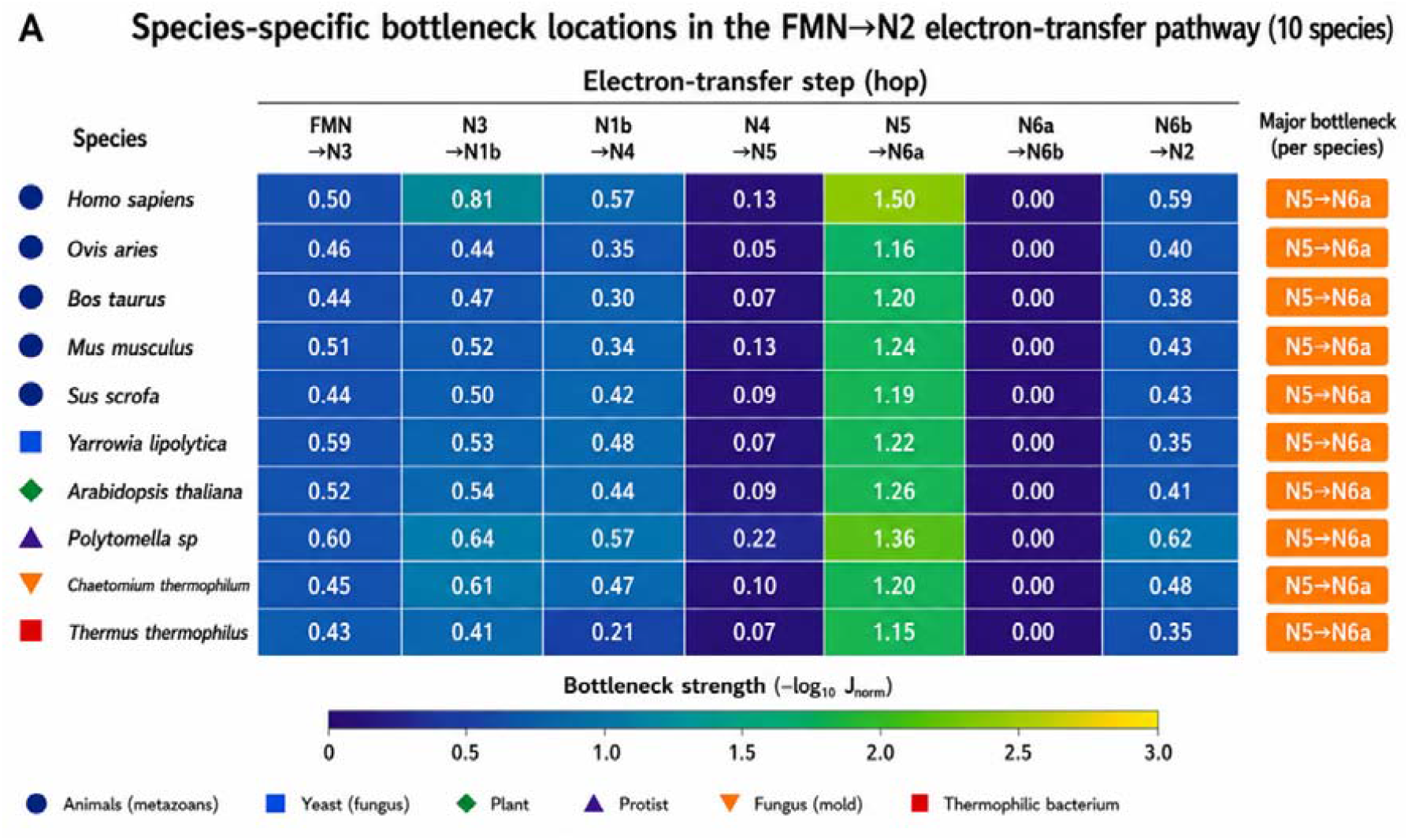
Species-specific bottleneck mapping in the FMN→N2 electron-transfer pathway. Heatmap showing bottleneck strength across individual electron-transfer steps in the mitochondrial complex I redox chain (FMN→N3→N1b→N4→N5→N6a→N6b→N2) for 10 species. Bottleneck strength is quantified as –log_10_(J_norm_), where J_norm_ represents the normalized electronic coupling between adjacent redox centers. Warmer colors indicate stronger kinetic bottlenecks (i.e., weaker electronic coupling and slower electron transfer), whereas cooler colors indicate more efficient transfer steps. Across all species analyzed, the N5→N6a step consistently exhibits the strongest bottleneck, identifying it as a conserved rate-limiting feature of the electron-transfer pathway. Despite this conserved bottleneck location, substantial variation in bottleneck strength is observed across species, indicating differences in electronic coupling at this step. The right panel summarizes the dominant bottleneck step for each species. Symbol annotations indicate phylogenetic groups: animals (metazoans), yeast (fungi), plant, protist, filamentous fungus (mold), and thermophilic bacterium.

Together, these results demonstrate that electron transport in complex I is governed by a structurally conserved kinetic bottleneck, establishing a bottleneck-controlled transport architecture in which a single spatially defined constraint dominates pathway efficiency.

### Electron transport dynamics are controlled by a single rate-limiting step

We next sought to determine how the structurally conserved bottleneck influences global transport dynamics. To this end, we simulated electron transfer along the redox chain using a master-equation-based transport model with structure-derived electronic couplings. Population dynamics were computed by solving for the time evolution of site populations across the redox network. The resulting population trajectories at the terminal site (N2) reveal pronounced variation in transport kinetics across species (**Fig. 2A**). While some systems exhibit rapid accumulation of population at N2, others display substantial delays, indicating marked differences in overall electron-transfer efficiency. To quantify these differences, we defined a dimensionless transport delay index, τ_50_, corresponding to the time required to reach 50% population at N2. Across species, τ_50_ spans a wide dynamic range (**Fig. 2B**), demonstrating that relatively small variations in microscopic coupling strengths are amplified into large differences in macroscopic transport times. To assess whether this variability arises from distributed contributions across the pathway or from a dominant step, we compared τ_50_ against the minimum electronic coupling (J_min_) for each species. We observe a strong scaling relationship between τ_50_ and J_min_ (**Fig. 2C**), indicating that global transport dynamics are tightly controlled by the weakest coupling in the system. Notably, this relationship persists despite substantial variation in non-limiting steps, indicating that contributions from faster transitions are effectively buffered and do not influence system-level kinetics. Systems with weaker bottleneck couplings consistently exhibit slower transport, demonstrating that variation in a single microscopic parameter is sufficient to reshape the entire transport profile. These results establish that electron transport in complex I operates in a bottleneck-dominated regime, in which system-level dynamics collapse onto a single rate-limiting step.

**Figure 2.**
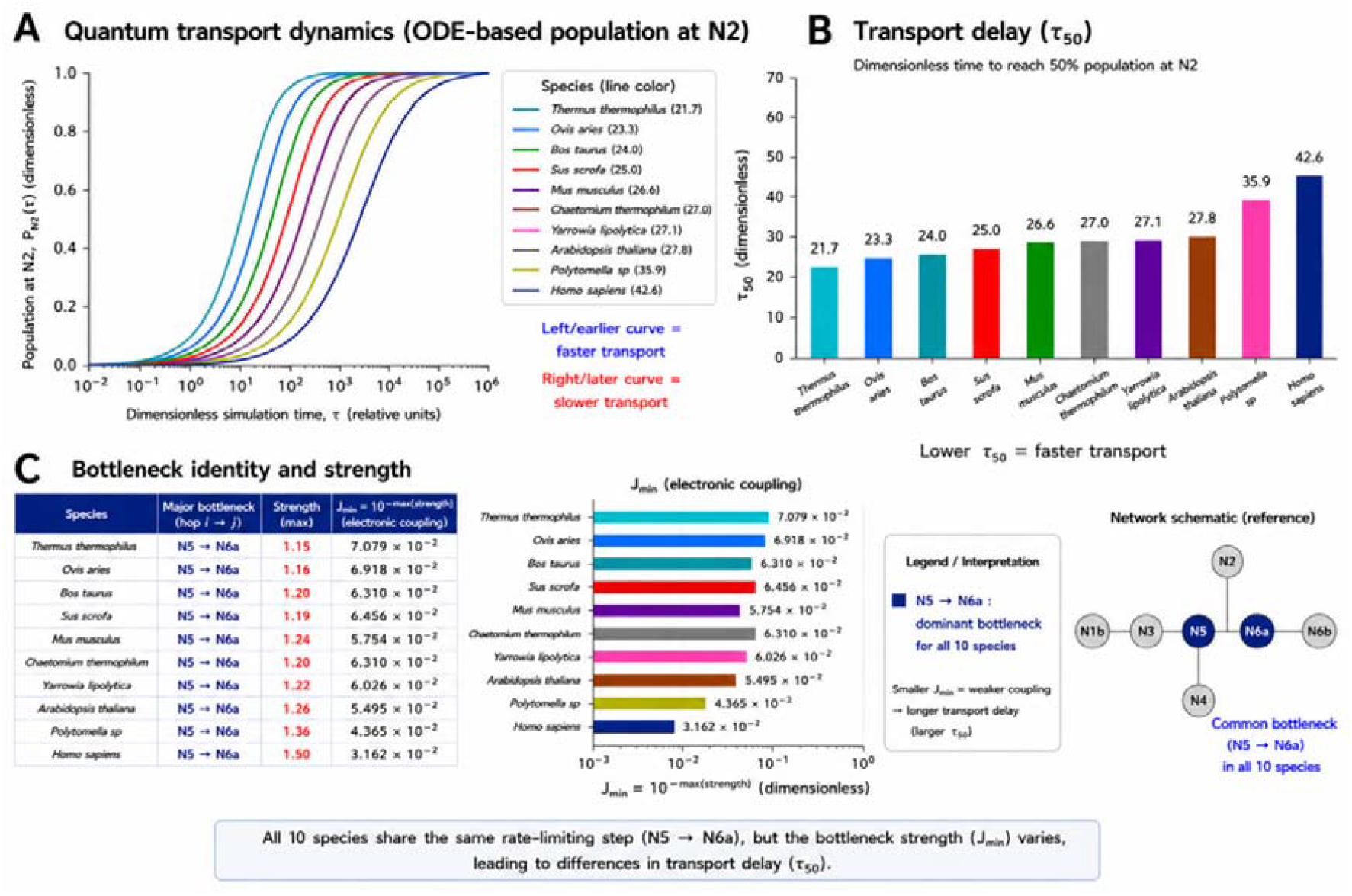
A conserved N5→N6a bottleneck controls species-specific electron transport dynamics. (A) ODE-based electron transport dynamics. Population transfer to the terminal acceptor (N2) as a function of dimensionless time (τ), computed from a sequential electron-transfer model parameterized by structure-derived coupling strengths. Curves are ordered by transport speed: earlier rise indicates faster transport. Substantial variation in kinetics is observed across species despite a conserved pathway architecture. (B) Transport delay across species. Characteristic transport time (τ50, time to reach 50% population at N2) is shown for each species. Lower τ50 corresponds to faster electron transfer. Transport delays span a broad dynamic range, indicating species-specific differences in pathway efficiency. (C) Bottleneck identity and coupling strength. All species share a common rate-limiting step at N5→N6a. However, the bottleneck strength (−log_10_ J_nor,_) varies across species, leading to differences in the effective coupling J_min_ = 10^−strength^. Stronger bottlenecks (lower J_min_) are associated with slower transport (higher τ50). Right: visualization of J_min_ across species. Inset: schematic of the electron-transfer network highlighting the conserved N5→N6a bottleneck.

### Geometric constraints, rather than local environment, determine bottleneck strength

To uncover the structural origin of the conserved kinetic bottleneck, we systematically examined two potential determinants: the local protein environment surrounding the redox centers and the geometric arrangement of the Fe–S clusters. We first quantified bottleneck strength at the N5→N6a transition across species using a structure-mapped coupling metric (**Fig. 3A**). Despite moderate variation in magnitude, all species exhibit a bottleneck at the same structural location, reinforcing that the rate-limiting step is not stochastic but embedded within a conserved architectural feature of complex I. We next tested whether local chemical environment contributes to variation in bottleneck strength by quantifying the fraction of polar residues within a defined radius around the N5–N6a region. No significant correlation was observed between local polarity and bottleneck strength (**Fig. 3B**; R^2^ = 0.19, *p* = 0.21), suggesting that local electrostatic environment is not the dominant determinant of variation in electron transfer efficiency. We note that, with ten species, the absence of a significant correlation should be interpreted with caution and does not exclude a contributing role for local environment; rather, it indicates that local chemical tuning alone is insufficient to account for the observed system-level transport limitation. In contrast, a dependence emerges when considering geometric factors. Bottleneck strength increases with the inter-cluster distance between N5 and N6a (**Fig. 3C**). While this relationship is relatively weak when considering all species together (R^2^ = 0.22, *p* = 0.18), a strikingly strong correlation is observed within eukaryotic systems (R^2^ = 0.95, *p* = 1.1 × 10^−5^), indicating that geometric separation is a dominant determinant of transport limitation in structurally conserved complexes. We restricted this correlation to eukaryotes a priori because the bacterial outlier (Thermus thermophilus) has a markedly shorter N5–N6a separation and a distinct subunit context, so that pooling it with the structurally conserved eukaryotic complexes mixes two regimes; the weaker all-species trend (R^2^ = 0.22) reflects this heterogeneity rather than an absence of a geometric dependence. Given the small number of species, we interpret the eukaryotic correlation as indicative rather than definitive.

**Figure 3.**
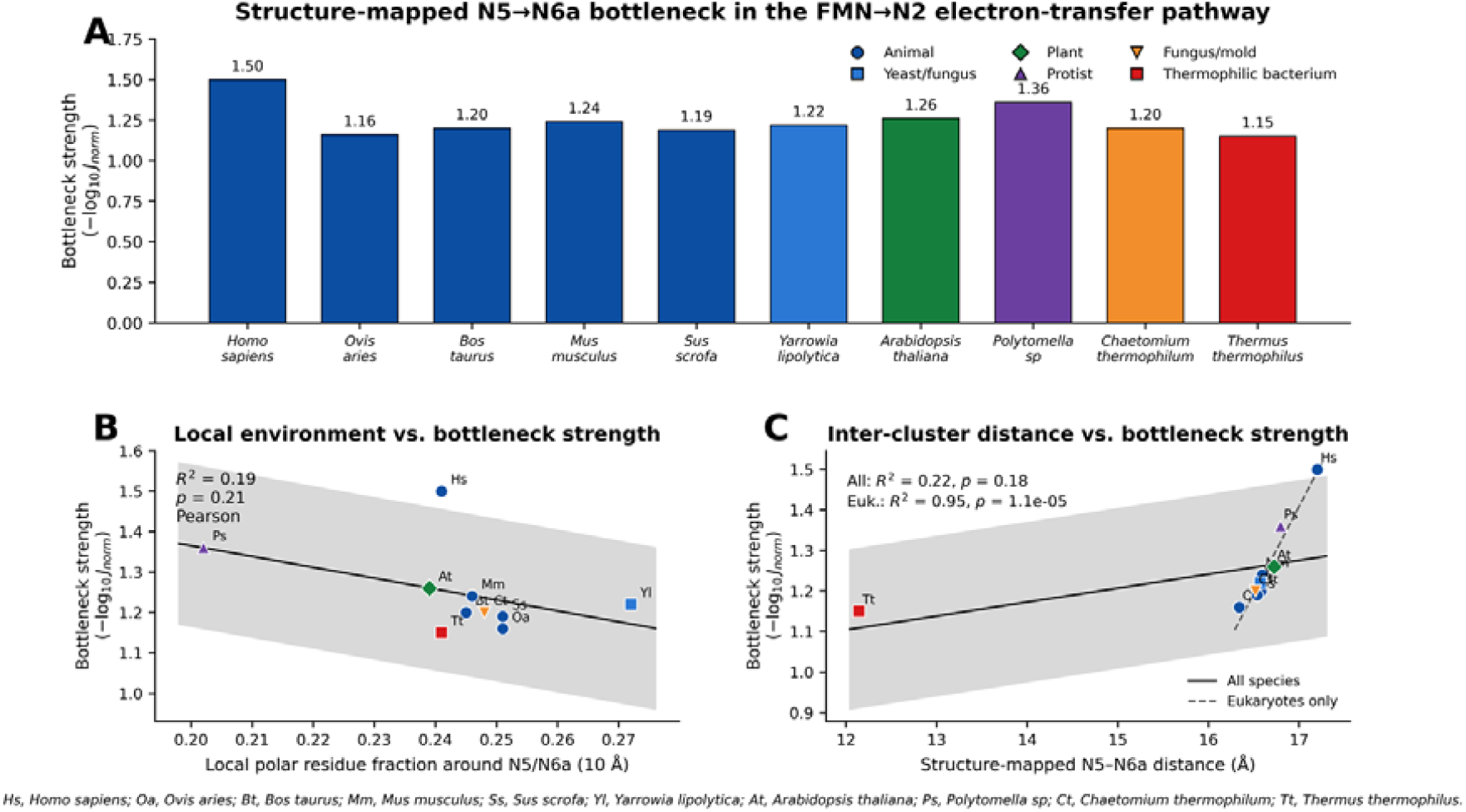
| Structural determinants of the N5→N6a bottleneck across species. (A) Structure-mapped bottleneck strength across species. Bottleneck strength (−log_10_ J_norm_) at the N5→N6a step is shown for all species. Despite conservation of the bottleneck location, substantial variation in strength is observed, indicating species-specific differences in electronic coupling. Colors denote phylogenetic groups. (B) Local environment does not strongly predict bottleneck strength. Bottleneck strength is plotted against the local polar residue fraction within 10 Å of the N5–N6a region. A weak and non-significant correlation (Pearson r, *p*-value shown) suggests that local chemical environment alone is insufficient to explain variations in coupling strength. (C) Structural distance governs bottleneck strength. Bottleneck strength is plotted against the centroid distance between N5 and N6a. A weak global trend contrasts with a strong correlation within eukaryotes, indicating that inter-cluster distance is a primary determinant of electronic coupling in conserved architectures. Solid line indicates fit for all species with 95% confidence interval (shaded), and dashed line shows the fit restricted to eukaryotes.

Importantly, this relationship is consistent with the exponential distance dependence predicted by electron tunneling theory, providing a direct physical basis for the observed scaling behavior. However, the emergence of a single dominant bottleneck is not a trivial consequence of this formulation. In a multi-step transport system, multiple weak links could in principle contribute comparably to overall transport limitation. Instead, our results reveal a strongly hierarchical regime in which heterogeneity in inter-cluster spacing produces a sharply defined weakest link that dominates system-level dynamics. To assess whether this conclusion depends on the specific definition of inter-cluster distance, we compared centroid-based distances with edge-to-edge corrected distances (**Supplementary Fig. S1**). Although distance correction systematically reduces the absolute N5–N6a separation and strengthens electronic coupling, the identity of the dominant bottleneck remains unchanged across all species. Consistent with this, the distance–strength relationship persists under both distance definitions (Supplementary Fig. S2). While the correlation strength and slope are reduced after correction (R^2^ decreases from ∼0.91 to ∼0.49), the positive dependence between distance and bottleneck strength remains intact, indicating that the observed trend is not an artifact of a specific geometric metric. These results demonstrate that the conserved bottleneck is robust to variations in structural measurement and arises from intrinsic spatial constraints within the protein architecture, as further summarized by structure-mapped determinants (Table 1). Accordingly, variation in bottleneck strength is governed primarily by geometric organization rather than local chemical environment, defining a structure-controlled transport regime in which the spatial arrangement of redox centers dictates electron transfer efficiency.

**Table 1.**
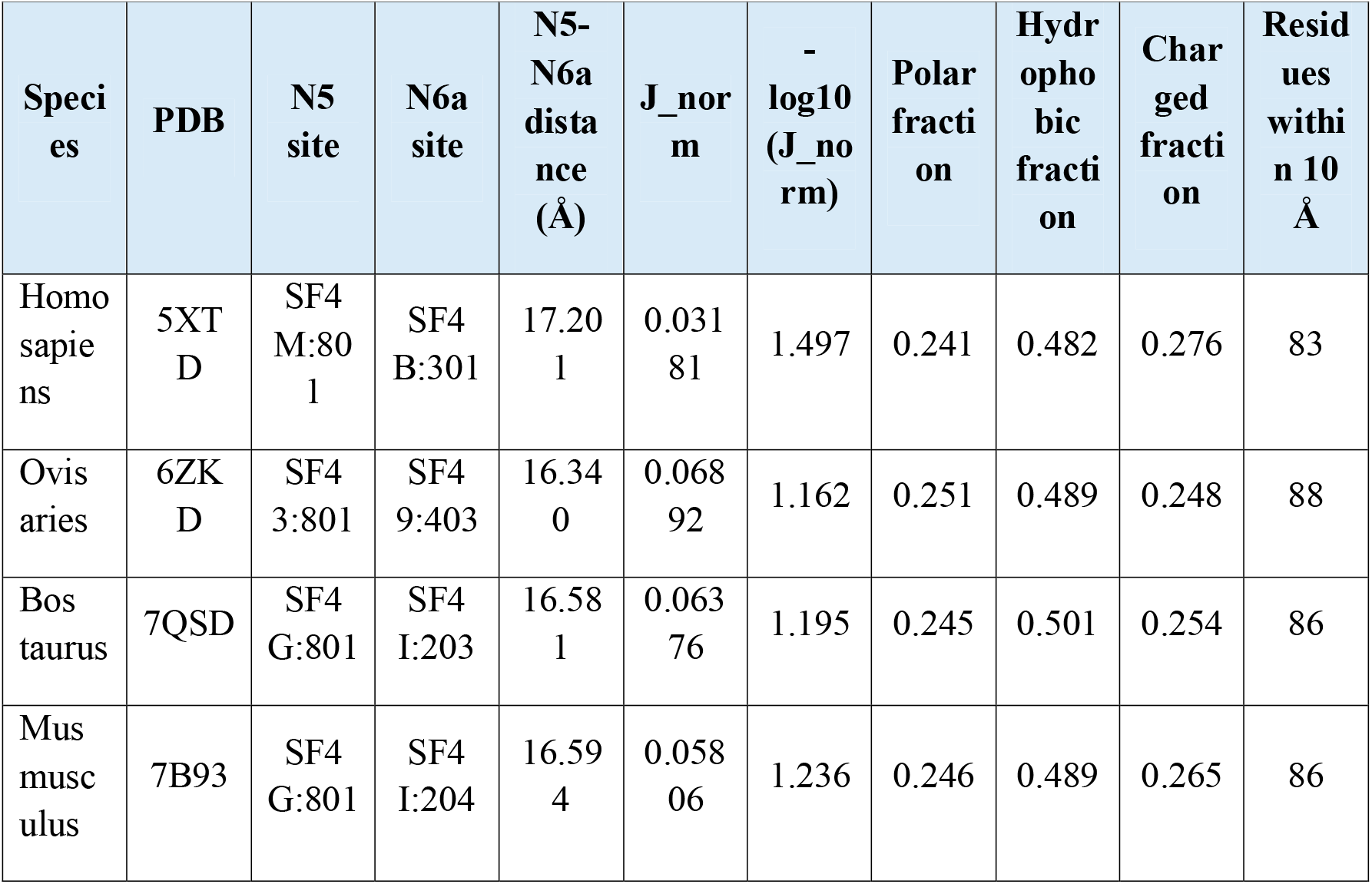
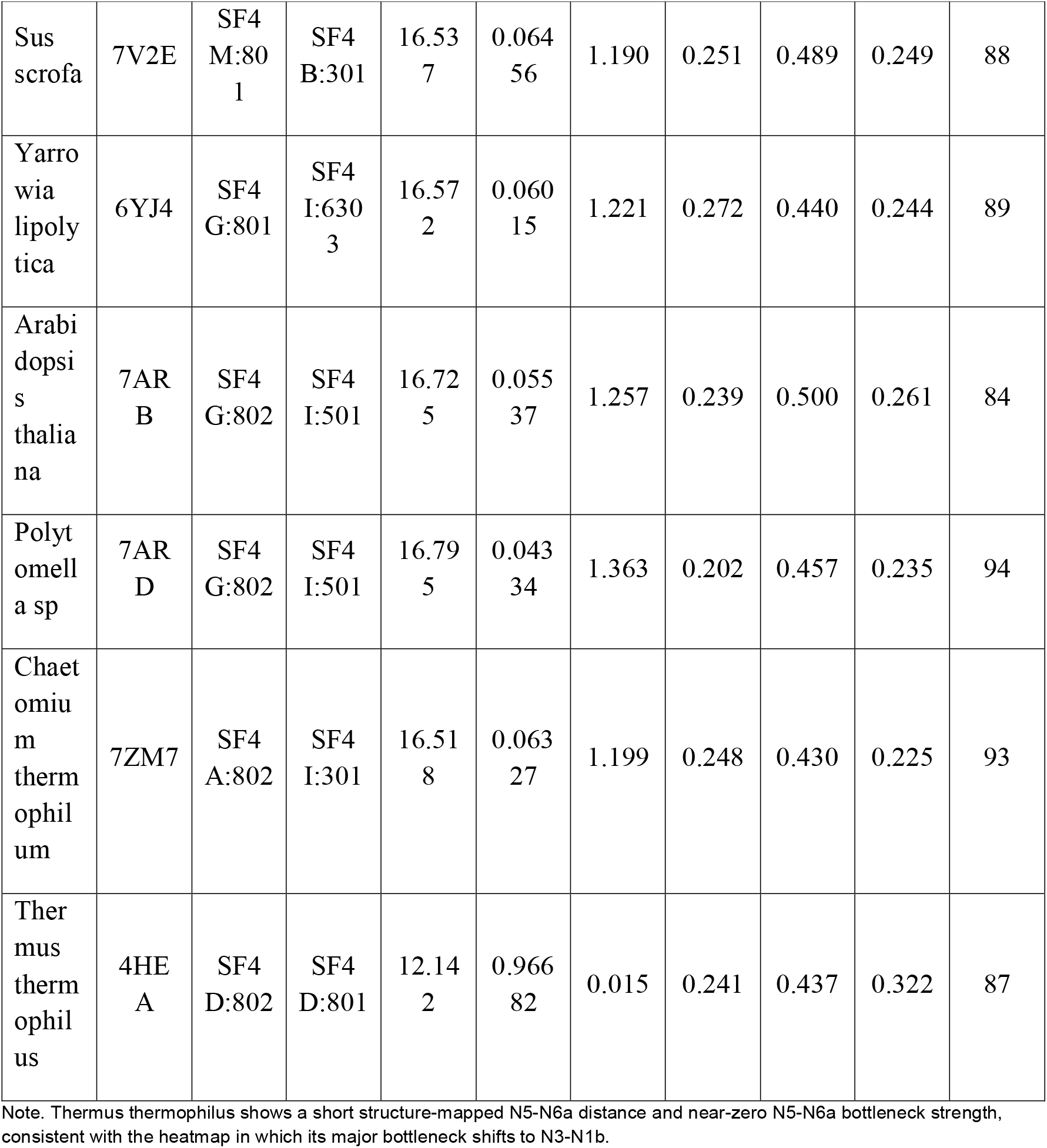
Structure-mapped N5-N6a bottleneck determinants. Values are taken directly from the structure-mapped N5-N6a calculation output. J_norm is the normalized distance-dependent coupling proxy used for the hop-wise bottleneck map.

### A structural cascade links protein architecture to transport efficiency

To integrate these observations into a unified framework, we examined how geometric variation at the bottleneck propagates across electronic and dynamical levels of electron transport (**Fig. 4**). Across species, the inter-cluster distance at the N5→N6a bottleneck spans a relatively narrow range (∼16–17 Å; **Fig. 4A**), yet even this modest variation is associated with measurable differences in bottleneck strength. In particular, increased separation between redox centers corresponds to stronger transport limitation, consistent with a geometry-dependent modulation of electronic coupling. To directly assess this relationship, we compared the minimum electronic coupling (J_min_) with effective transport rates (k_eff_ = 1/ τ_50_). A strong monotonic relationship is observed (Spearman ρ = 0.94, p = 6.7 × 10^−5^; **Fig. 4B**), indicating that variation in coupling at the bottleneck is directly translated into differences in system-level transport rates. Systems with weaker coupling exhibit consistently slower transport, demonstrating that microscopic electronic constraints propagate to macroscopic dynamics. Consistent with this, species-specific transport rates span a broad range **(Fig. 4C**), with slower systems corresponding to larger bottleneck distances and weaker coupling strengths. These results indicate that relatively small structural perturbations at the angstrom scale can give rise to substantial variation in transport efficiency. Together, these observations define a mechanistic cascade linking protein structure to function **(Fig. 4D**): geometric expansion at the bottleneck reduces electronic coupling, which in turn lowers transport rates and ultimately decreases pathway efficiency. In this framework, the N5–N6a interface emerges as a structurally encoded control point, where small geometric differences are amplified into system-level functional variation. The consistency of this cascade across diverse species suggests that bottleneck-mediated control of electron transport represents a general principle, in which protein architecture governs biological function through a hierarchy of coupled structural and dynamical constraints.

**Figure 4.**
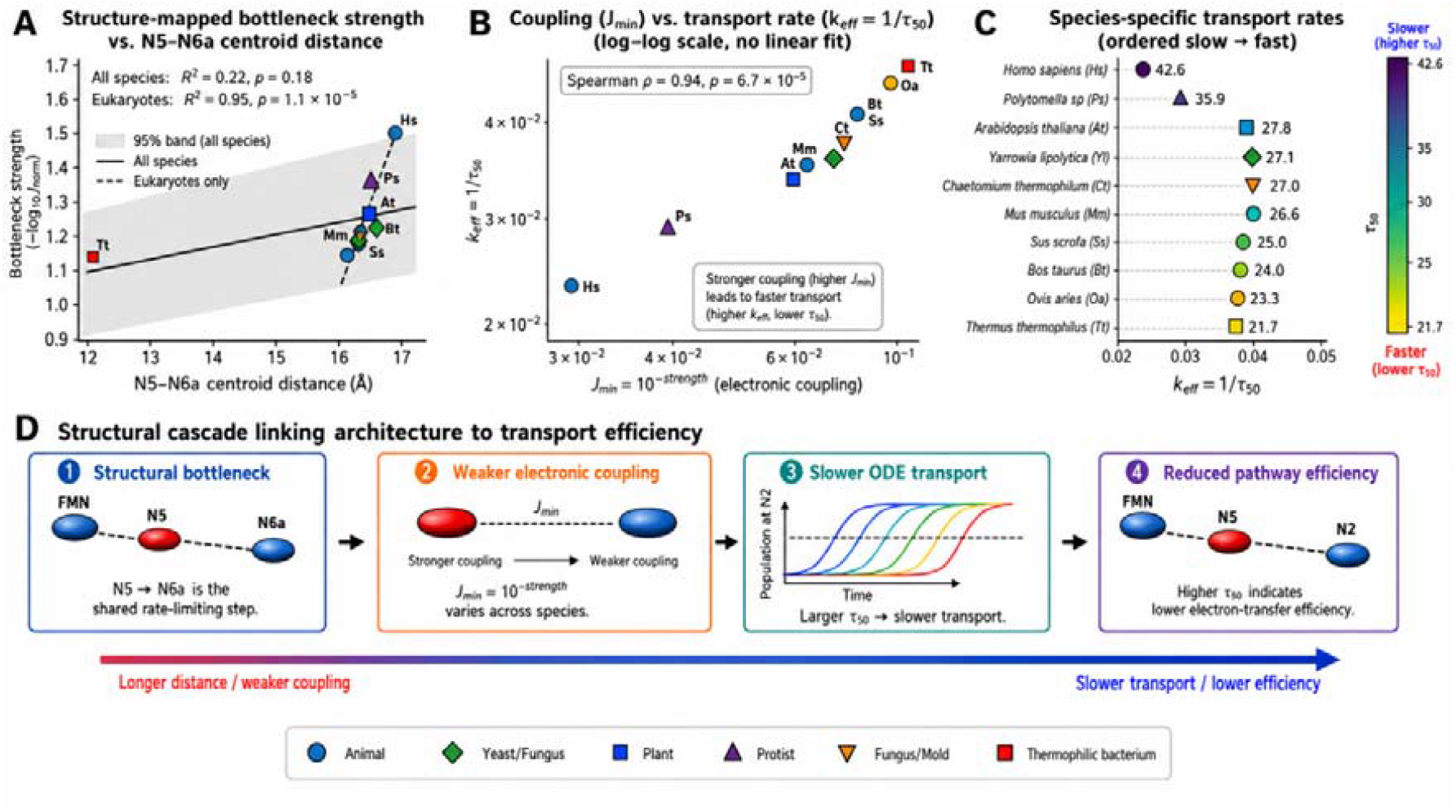
Structural control of electron transport efficiency via a conserved N5→N6a bottleneck. (A) Structure–coupling relationship at the bottleneck. Bottleneck strength (−log_50_ J_norm_) at the N5→N6a step is plotted against the corresponding centroid distance across species. A weak global trend contrasts with a strong correlation within eukaryotes, indicating that structural variation modulates electronic coupling in a lineage-dependent manner. (B) Coupling determines transport kinetics. Effective transport rate (k_eff_ = 1/τ50) scales with bottleneck coupling (J_min_) on a log–log axis. Stronger coupling yields faster transport, demonstrating that a single rate-limiting step quantitatively governs pathway kinetics across species (Spearman ρ shown). (C) Species-resolved transport efficiency. Species-specific transport rates (k_eff_) are ordered from slow to fast. Despite a conserved bottleneck identity (N5→N6a), variations in coupling strength give rise to a wide dynamic range of transport efficiencies. (D) Structural cascade linking architecture to function. Schematic illustrating the mechanistic cascade: increased N5–N6a distance weakens electronic coupling, delays population transfer, and reduces overall electron transport efficiency. This establishes a direct mapping from structural geometry to functional output.

### Decoherence modulates quantum transport while preserving bottleneck-limited dynamics

To investigate how environmental interactions influence electron transport, we incorporated decoherence into a continuous-time quantum-walk framework as an effective system– environment coupling parameter (γ) and analyzed its impact across species. The dependence of transport on decoherence strength reveals a non-monotonic behavior (**Fig. 5A**). As γ increases from the coherent limit, population transfer to the terminal site (N2) initially improves, reaches a maximum, and subsequently decreases at higher decoherence levels. This behavior is consistent with environment-assisted quantum transport, in which intermediate decoherence enhances transport efficiency while excessive dephasing suppresses coherent propagation. Importantly, the position and magnitude of this optimal regime vary across species. Systems with larger N5–N6a distances (corresponding to weaker bottleneck coupling) exhibit reduced peak transfer and a narrower optimal regime, indicating increased sensitivity to decoherence. To quantify this effect, we defined a decoherence sensitivity metric based on the response of terminal population to changes in γ. Sensitivity shows a positive trend with bottleneck distance (**Fig. 5B**; Pearson r = 0.59, *p* = 0.073), consistent with a tendency for systems with greater geometric separation to be more susceptible to environmental perturbations. This correlation does not reach statistical significance at the conventional threshold (p = 0.073, n = 10) and should therefore be regarded as a suggestive trend rather than an established relationship. We next examined representative transport dynamics under weak and strong decoherence regimes (**Fig. 5C**). Under weak decoherence, population transfer proceeds more efficiently, whereas strong decoherence suppresses transport and leads to slower accumulation at the terminal site. Despite these environment-dependent effects, the relative ordering of species is preserved: systems with weaker bottleneck coupling consistently exhibit slower transport across all decoherence regimes. This indicates that environmental interactions modulate transport efficiency but do not override the structural constraint imposed by the bottleneck. These results demonstrate that electron transport in complex I is governed by an interplay between structure and environment, in which geometric constraints define a bottleneck-limited baseline while decoherence modulates transport efficiency within that framework.

**Figure 5.**
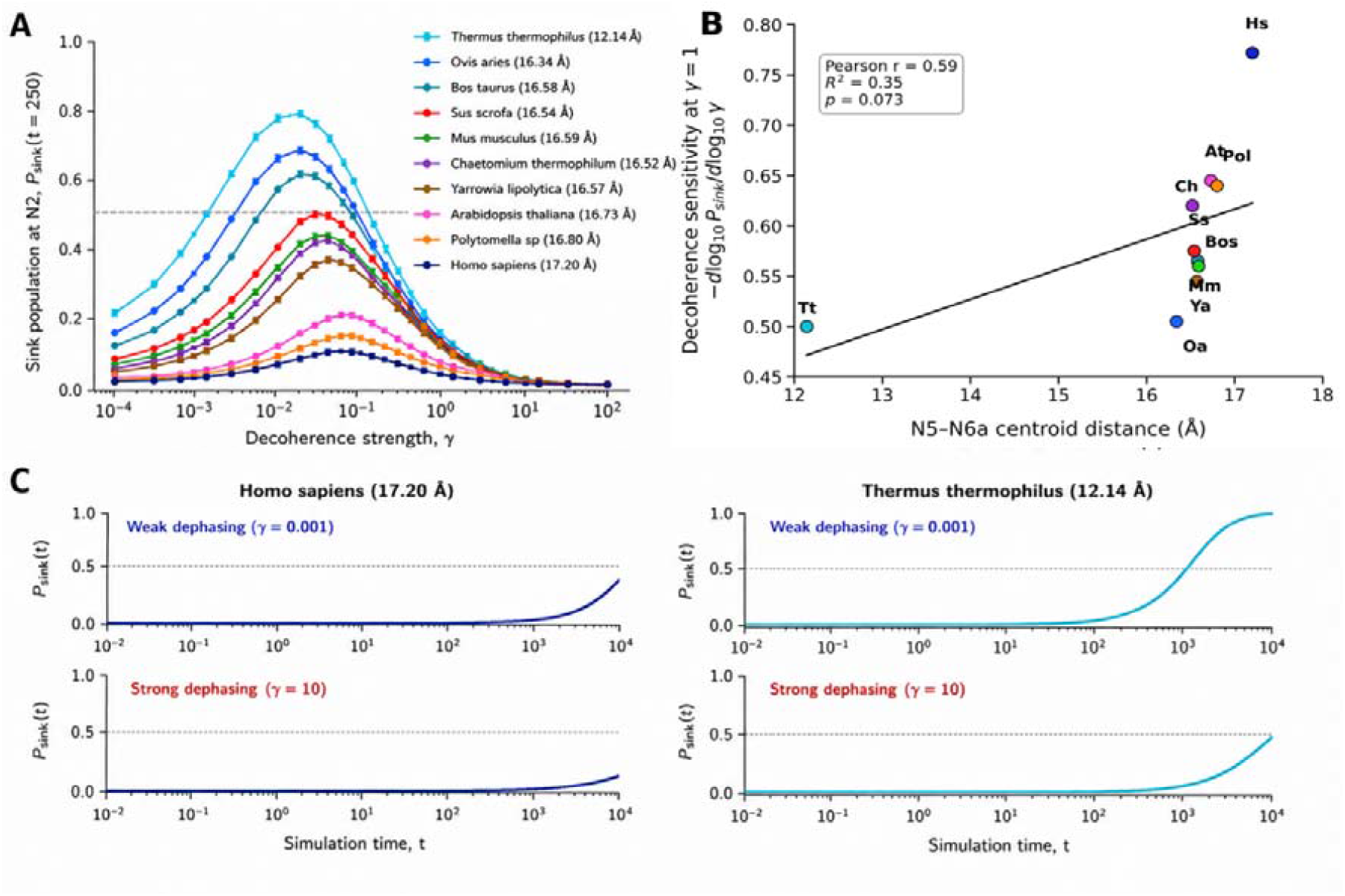
Sink-including Lindblad dynamics reveal decoherence-dependent electron transport bottlenecks. (A)Decoherence-dependent sink population across species. Population transferred to the terminal acceptor (N2 sink) as a function of decoherence strength (γ), computed using a Lindblad master equation including site-local dephasing and irreversible trapping at N2. Each curve corresponds to one species parameterized by its structure-derived N5–N6a centroid distance. Intermediate decoherence maximizes transport efficiency (environment-assisted transport), whereas both low and high γ reduce sink population. Species with shorter N5–N6a distances (stronger coupling) exhibit higher peak transfer. (B) Sensitivity of transport to decoherence vs. structural distance. Decoherence sensitivity, defined as evaluated at γ = 1, plotted against N5–N6a centroid distance. Each point represents one species (n = 10). Color indicates structural distance (Å). A positive correlation indicates that longer distances (weaker electronic coupling) lead to stronger dependence on environmental decoherence. Spearman correlation coefficient (ρ) and p-value are shown. (C) Representative sink-trapping trajectories under weak and strong dephasing. Time evolution of sink population Psink(t) for representative species: • Homo sapiens (long distance, weaker coupling) • Thermus thermophilus (short distance, stronger coupling) Top panels: weak dephasing (γ = 10^−3^), Bottom panels: strong dephasing (γ = 10^1^) Under weak dephasing, coherent transport enhances transfer efficiency, particularly in strongly coupled systems. Under strong dephasing, transport becomes diffusion-limited and overall efficiency decreases.

### Quantum-walk analysis reveals residue-mediated alternative electron-transfer pathways

To determine whether electron transport in complex I is confined to a single canonical pathway or can be redistributed across the network, we analyzed pathway-level transport using a continuous-time quantum-walk framework. We first identified the dominant residue-mediated alternative pathways connecting N5 to N6a across species (**Fig. 6A**). These pathways are consistently mediated by short chains of amino acid residues, forming indirect connections that bypass the canonical direct transition. Despite variation in residue identity, the presence of such alternative routes is conserved, indicating that the redox network supports multiple structurally encoded pathways. To quantify the contribution of these routes, we computed pathway-resolved transport weights using quantum-walk flux. Across all species, alternative residue-mediated pathways exhibit substantial flux relative to the direct N5→N6a transition (**Fig. 6B**). In several cases, the cumulative flux through alternative pathways exceeds that of the direct route, indicating that, within this model, electron transport is not restricted to a single dominant path but can be distributed across a network of competing channels. We verified that this flux redistribution persists across a range of bridge on-site energy offsets and direct-coupling scaling, so that the dominance of alternative routes is not an artifact of a single parameter choice.

**Figure 6.**
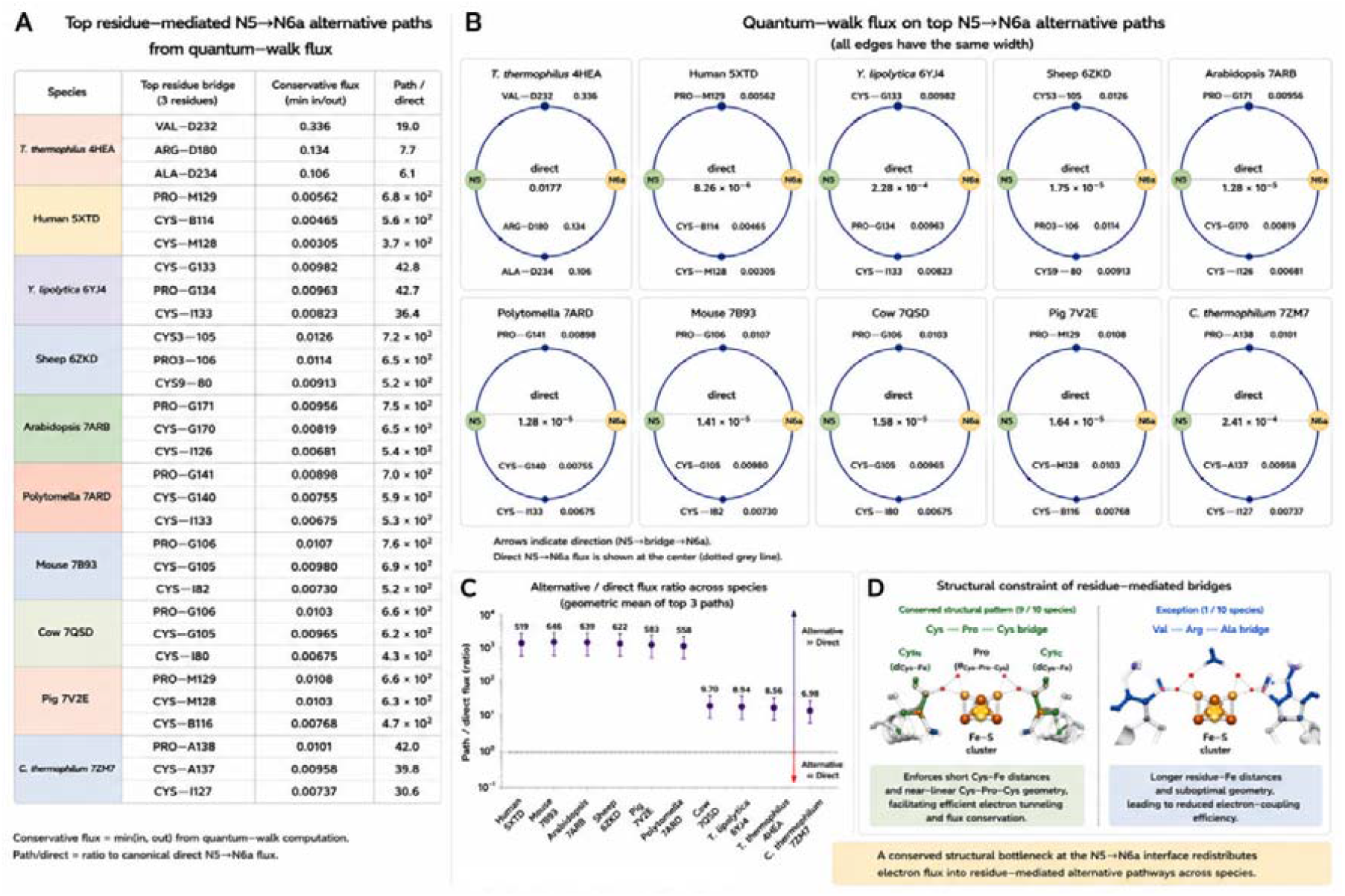
Residue-mediated alternative electron transfer pathways at the N5→N6a interface revealed by quantum-walk flux analysis. (A) Top residue-mediated alternative pathways connecting N5 to N6a across representative species, ranked by conservative quantum-walk flux (defined as min[in, out]). For each species, the top three residue bridges are shown with their corresponding flux values and relative path strength normalized to the canonical direct N5→N6a transfer (Path/direct). (B) Circular network representation of quantum-walk flux distribution for the top N5→N6a alternative pathways. All edges are drawn with equal width to isolate flux magnitude (annotated numerically) from visual bias. Arrows indicate the direction of electron flow (N5 → residue bridge → N6a), while the central dashed line denotes the direct N5→N6a flux. (C) Ratio of alternative-to-direct flux across species, computed as the geometric mean of the top three residue-mediated pathways per species. Values >1 indicate dominance of alternative pathways over the direct route, highlighting a conserved redistribution of electron flux. Error bars represent variability among the top-ranked paths. (D) Structural basis of residue-mediated electron transfer. A conserved Cys–Pro–Cys motif (observed in 9/10 species) enforces short Cys–Fe distances and near-linear geometry, facilitating efficient electron tunneling. In contrast, a rare Val–Arg– Ala configuration (1/10 species) exhibits longer distances and suboptimal geometry, consistent with reduced coupling efficiency. A conserved structural bottleneck at the N5–N6a interface redirects electron flow into residue-mediated alternative pathways across species, indicating a universal design principle in Complex I electron transport.

This redistribution is further supported by the ratio of alternative to direct pathway flux (**Fig. 6C**). Across species, alternative pathways consistently contribute at comparable or greater levels than the canonical transition, indicating that the bottleneck at N5–N6a does not simply suppress transport but instead redirects electron flow into parallel routes. At the structural level, these alternative pathways are enabled by conserved residue-mediated bridging motifs (**Fig. 6D**). In most species, cysteine–proline–cysteine configurations form geometrically favorable connections that maintain short effective tunneling distances and facilitate electron transfer. In contrast, deviations from this motif are associated with longer distances and reduced coupling efficiency, highlighting the structural constraints that govern pathway accessibility. Together, these results demonstrate that electron transport in complex I is not limited to a single linear pathway but instead operates as a network process, in which a structurally encoded bottleneck actively redistributes electron flux into residue-mediated alternative routes. This reveals an additional layer of organization in biological electron transport, where protein architecture governs not only the efficiency but also the topology of electron flow.

## Discussion

Electron transport in mitochondrial complex I has traditionally been interpreted within classical hopping frameworks, in which electrons propagate sequentially through localized redox centers governed by stochastic rate processes. ^4 10^ While such models capture aspects of overall kinetics, they do not account for the quantum nature of electron motion at the nanoscale, where coherence, interference, and environmental interactions can fundamentally influence transport behavior. ^20 21^ Here, we reformulate electron transport as a continuous-time quantum walk on a structure-derived redox network, providing a unified framework that integrates coherent dynamics, environmental decoherence, and network-level connectivity.^17^ Within this framework, we identify a conserved structural bottleneck at the N5–N6a interface that governs transport dynamics across species. ^22^ Importantly, this bottleneck is not merely a kinetic feature but emerges directly from the underlying Hamiltonian as a suppression of electronic coupling, providing a fundamentally different physical interpretation of rate limitation.^23^ We note, however, that the primary bottleneck identified here is largely geometry-driven and can therefore emerge in both incoherent hopping and coherent transport descriptions. The principal contribution of the continuous-time quantum-walk framework is not the existence of the bottleneck itself, but the ability to resolve pathway-level flux redistribution, competing transport channels, and decoherence-dependent network dynamics within a unified Hamiltonian formalism.

A central result of this study is that electron transport operates in a bottleneck-dominated regime, in which system-level dynamics collapse onto a single weakly coupled transition.

In the quantum description, this bottleneck acts as a barrier to amplitude propagation, limiting transmission of the wavefunction across the network. Unlike classical sequential transport, where populations move stepwise, quantum transport depends on coherent amplitude propagation, making the system intrinsically sensitive to a single weak link. This establishes a direct mapping between microscopic coupling and macroscopic transport behavior. We further demonstrate that the origin of this bottleneck is geometric rather than chemical. Coupling strength is primarily determined by inter-cluster distance, with negligible contribution from local protein environment. This indicates that protein architecture defines the Hamiltonian governing electron transport, with spatial arrangement directly encoding network topology and coupling strengths. Importantly, this structure–function mapping is robust to variations in structural metrics, confirming that the bottleneck reflects intrinsic spatial constraints rather than modeling artifacts. Incorporating environmental decoherence reveals that transport operates in a regime where coherent and incoherent processes coexist. Transport efficiency exhibits a non-monotonic dependence on decoherence strength, consistent with environment-assisted quantum transport. Moderate decoherence enhances transport by suppressing destructive interference, whereas strong decoherence leads to diffusive behavior and reduced efficiency.

Notably, however, the relative ordering of transport rates across species is preserved, indicating that environmental effects modulate, but do not override, the structural constraint imposed by the bottleneck. Beyond its effect on efficiency, the bottleneck also reshapes how electron flow is distributed across the network. Quantum-walk flux analysis reveals that the N5–N6a bottleneck does not simply suppress transport along the canonical pathway, but actively redistributes electron flow into residue-mediated alternative pathways. These alternative routes, formed by short chains of amino acids, carry substantial flux and in several cases exceed the direct pathway contribution. This demonstrates that electron transport in complex I is inherently a network process, rather than a strictly linear sequence of steps. This observation extends the concept of rate limitation: the bottleneck does not merely slow transport, but organizes it. Within the quantum framework, suppression of amplitude at the bottleneck enhances exploration of parallel pathways, leading to redistribution of probability flux across the network. This provides a mechanistic basis for pathway-level flexibility in biological electron transport systems. The conservation of the N5–N6a bottleneck across species suggests that this feature serves a functional role. A structurally encoded bottleneck may regulate electron flux, preventing excessive charge accumulation and mitigating deleterious side reactions, while the presence of alternative pathways provides redundancy and robustness. This combination of constraint and flexibility suggests an evolutionarily optimized design in which geometry defines a stable baseline and network connectivity enables adaptive redistribution of transport. Together, these findings support a working framework for structurally encoded control of biological electron transport, in which transport efficiency in multi-step redox systems is governed by a structurally encoded bottleneck that arises from geometric constraints, persists under environmental decoherence, and actively redistributes electron flow across alternative pathways. In this framework, protein architecture encodes not only the efficiency but also the topology of electron transport through a hierarchy of geometric constraints, quantum dynamics, and network-level connectivity. This study focuses on mitochondrial complex I; whether the same bottleneck- and-redistribution behavior extends to other biological electron-transfer systems, such as photosynthetic complexes and redox enzymes, is an open question that will require dedicated analysis of those architectures. More broadly, these results highlight the importance of integrating structural biology with quantum transport theory, placing biological electron transport within the emerging framework of quantum biology.

## Methods

### Conceptual framework for structure-driven quantum transport

Electron transport in mitochondrial complex I was modeled as a structure-constrained quantum dynamical process occurring along a discrete network of redox centers. ^24^ Unlike conventional classical hopping models, which describe transport as stochastic population transfer between localized states, the present framework treats electron motion as a coherent quantum process governed by wavefunction propagation and its interaction with the surrounding environment. ^25 17^

The central objective of this modeling framework is to disentangle the relative contributions of intrinsic structural geometry and environmental effects in governing transport efficiency. To this end, we establish a hierarchical formalism in which protein structural organization defines geometric constraints, which in turn modulate electronic coupling, thereby shaping the ensuing quantum dynamics and ultimately determining transport efficiency. This formulation enables direct interrogation of whether electron transport is primarily governed by local biochemical environment or by global structural organization, a question that cannot be resolved within purely classical frameworks. ^21,26^ Importantly, the quantum-walk formalism provides a natural description of transport in systems where coherence, interference, and environmental decoherence may coexist, making it particularly suitable for biological electron-transfer networks.

### Structural dataset and definition of redox centres

Cryo–electron microscopy (cryo-EM) structures of mitochondrial (and bacterial) Complex I were collected to construct a cross-species structural dataset spanning the evolutionary spectrum from bacteria to mammals.^27-29^ The analysed structures included representatives from Homo sapiens (PDB: 5XTD), Bos taurus (7QSD), Mus musculus (7B93), Ovis aries (6ZKD), Sus scrofa (7V2E), Arabidopsis thaliana (7ARB), Polytomella sp. (7ARD), Drosophila melanogaster (7P1L), Chaetomium thermophilum (7ZM7), Yarrowia lipolytica (6YJ4), Escherichia coli (7NYT), and Thermus thermophilus (4HEA). These species encompass wide variations in metabolic strategy, cellular environment, and evolutionary lineage, while preserving the conserved architecture of the canonical electron-transfer chain linking NADH oxidation to quinone reduction ^30^. Across all structures ^31^, the redox-active cofactors forming the primary electron-transfer pathway were systematically identified using ligand annotations provided in the deposited coordinate files. The analysed centres included flavin mononucleotide (FMN) and the iron–sulfur clusters N3, N1b, N4, N5, N6a, N6b, and N2. To enable quantitative structural comparison across species, each redox centre was represented using a geometry-based centroid definition. Fe–S clusters were represented by the centroid of the iron atoms composing each cluster, capturing the effective electronic centre of the delocalized redox-active unit while minimizing sensitivity to coordinating cysteine geometries. FMN was represented by the centroid of heavy atoms within the isoalloxazine ring system, corresponding to the π-electron-active region responsible for redox chemistry. All centroid coordinates were derived directly from experimentally determined atomic positions in the deposited PDB files without structural relaxation or modelling, thereby preserving native structural geometry ^32^. This enabled consistent cross-species mapping of the FMN → Fe–S electron-transfer chain for downstream analysis of spatial organization and electron transport topology.

### Cross-species node alignment

Cross-species structural comparison required consistent identification of equivalent Fe–S clusters despite substantial sequence divergence and species-specific insertions/deletions. To establish node correspondence in a structure-driven manner, each Fe–S cluster was represented by its Fe-atom centroid and assigned a local protein environment defined by residues proximal to the centroid. For each cluster centroid, neighbouring residues were collected using a spherical radial cutoff *r* (Å), defined by the minimum heavy-atom distance between any atom of a residue and the cluster centroid. The resulting set of residues was converted into a coarse-grained physicochemical description by classifying amino acids into three groups: charged (Asp, Glu, Lys, Arg, His), polar (Ser, Thr, Asn, Gln, Tyr, Cys), and hydrophobic (Ala, Val, Leu, Ile, Met, Phe, Trp, Pro, Gly). For each cluster, an environment fingerprint was then defined as the fractional composition of these classes within the local neighbourhood:

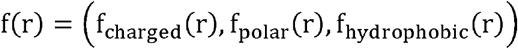

where each component corresponds to the fraction of residues belonging to the respective class and ∑ *f* = 1. Clusters were matched across species by minimizing the difference between fingerprints, enabling assignment of homologous redox nodes without requiring strict sequence alignment. For each species, Fe–S clusters were paired to the reference set by solving a minimum-distance matching problem in fingerprint space (using the human Complex I structure, 5XTD, as the reference), with the fingerprint dissimilarity quantified as the Euclidean distance ∥*f*(*a*)(*r*)− *f*(*b*)(*r*)∥. This procedure yielded a one-to-one correspondence between clusters that prioritizes conservation of the physicochemical microenvironment surrounding each redox centre, which is expected to be more evolutionarily constrained than primary sequence alone. To assess robustness of node assignment with respect to neighbourhood definition, matching was repeated across radial cutoffs spanning *r* =8–12 Å. Consistent assignments across this range were considered robust, while discrepancies were flagged for manual inspection in the context of the overall FMN → N2 chain topology and spatial ordering of redox centres. This environment-fingerprint alignment ^33^ provided a reproducible and species-agnostic procedure for mapping equivalent Fe–S nodes, supporting quantitative comparison of inter-centre distances and electron-transfer organization across the ten-species Complex I dataset.

### Local structural environment analysis

To examine whether bottleneck nodes are associated with distinct local protein environments, the structural neighbourhood surrounding each redox centre was systematically characterized. For each node, residues located within a 10 Å radial distance of the cluster centroid were identified based on minimum heavy-atom proximity. These residues were classified according to physicochemical properties, enabling quantification of the local microenvironment surrounding each redox site ^34^. For each node, the fractional composition of polar and hydrophobic residues within this neighbourhood was calculated. To assess environment-specific signatures associated with transfer constraints, polar enrichment and hydrophobic depletion were evaluated relative to the distribution observed for non-bottleneck nodes within the same species. Enrichment metrics were defined as deviations in residue-class fractions between bottleneck and non-bottleneck environments, allowing identification of systematic shifts in local polarity or hydrophobicity associated with structurally limiting electron-transfer steps ^35^. This analysis provided a consistent framework for linking local protein composition to bottleneck formation across species.

### Topological reconstruction of the redox-chain network

The electron-transfer pathway was reconstructed as a one-dimensional ordered network connecting FMN to the terminal N2 cluster. This was achieved using a nearest-neighbor algorithm based on Euclidean distance:

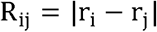

Starting from FMN, the next redox center was selected as the cluster minimizing R_ij_, and this procedure was iterated to produce the sequence:

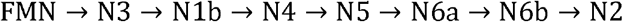

While this algorithm provides a geometry-driven reconstruction, the resulting topology was cross-validated against known biochemical assignments to ensure correct identification of functional redox sites. ^10^ This step ensures that the quantum-walk model is defined on a physically meaningful network rather than an arbitrary graph. ^20^

### Structural representation and coarse-graining of redox centers

Atomic-resolution structures of complex I were coarse-grained into a set of discrete redox sites corresponding to FMN and Fe–S clusters.^10^ Because electron transfer occurs through spatially extended orbitals rather than individual atoms, each redox center was represented by a single effective coordinate.

For Fe–S clusters, the coordinate was defined as the centroid of Fe atoms: ^3^

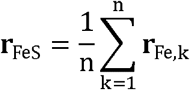

reflecting the dominant role of iron atoms in mediating electronic coupling.^20^ This choice reduces sensitivity to ligand geometry and preserves the physically relevant tunneling center. ^36^

For FMN, the centroid was computed over heavy atoms:

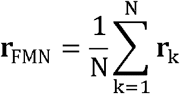

which ensures robustness against hydrogen placement and protonation-state ambiguity.^37^ This coarse-graining step defines the nodes of the quantum network and represents a balance between atomistic accuracy and computational tractability. ^38^

### Structural definition of redox centers and distance metrics

Redox centers in mitochondrial complex I were represented using a coarse-grained structural model. For Fe–S clusters, the effective coordinate was defined as the centroid of Fe atoms:

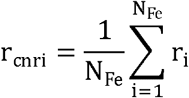

where r_i_ denotes the Cartesian coordinates of individual Fe atoms.

For flavin mononucleotide (FMN), the centroid was computed over heavy atoms. Inter-center distances were calculated using two distinct metrics: ^3^

1. Centroid–centroid distance

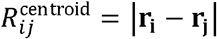
2. Edge-to-edge distance

To approximate the physically relevant tunneling distance, an edge-corrected distance was defined as:

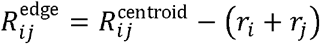

where r_i_ and r_j_ represent the effective radii of the corresponding redox centers, estimated from the maximal radial extent of Fe atoms within each cluster. This correction accounts for the finite spatial extent of cofactors and approximates the donor–acceptor edge separation relevant for electron tunneling. ^39^

### Distance-dependent electronic coupling

Electronic coupling between adjacent redox centers was modeled as an exponential function of inter-center distance, consistent with electron tunneling theory: ^40^

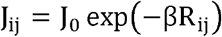

where β is the decay constant and R_ij_ is either centroid-based or edge-corrected distance. ^18 41^ To compare relative coupling strengths across species, couplings were normalized within each system: ^3^

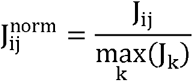

and expressed as:

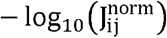

which emphasizes weak-coupling (bottleneck) regimes. ^10^

### Identification of bottlenecks

To identify structurally induced constraints on electron transfer, hop-wise perturbations were computed for each species along the canonical FMN → N2 pathway. For each donor– acceptor step *i*, the relative perturbation in transfer efficiency was quantified using the logarithmic distance-based formulation derived from the Moser–Dutton framework:

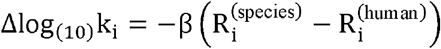

where 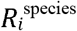 and 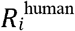 denote the inter-centre distances for hop *i* in the analysed species and in Homo sapiens (5XTD), respectively. The dominant bottleneck was defined as the hop exhibiting the greatest predicted reduction in transfer efficiency:

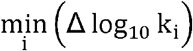

This hop corresponds to the step most strongly limiting electron-transfer efficiency relative to the human reference. To evaluate pathway-level effects, cumulative perturbation across the entire FMN → N2 chain was computed as:

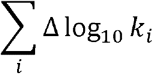

This quantity represents the net structural modulation of transfer efficiency across the multi-step redox relay ^40^. The contribution of the dominant hop to the overall pathway perturbation was then assessed by comparing its magnitude to the cumulative perturbation, thereby quantifying bottleneck dominance. This metric distinguishes between systems in which transfer efficiency is limited by a single structurally constrained step and those in which perturbations are distributed across multiple hops. This formulation enables topology-aware comparison of electron-transfer constraints across evolutionarily diverse Complex I architectures.

### Electron-transfer distance calculation

Electron-transfer efficiency along the FMN → Fe–S relay was evaluated using the distance-dependent tunnelling framework introduced by Moser and Dutton ^18^, which relates biological electron-transfer rates to donor–acceptor separation through an exponential decay formalism. In this model, the rate constant is governed primarily by the edge-to-edge distance between redox centres:

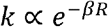

where *R* denotes the donor–acceptor distance and *β* is the tunnelling attenuation factor describing exponential decay of electronic coupling through the protein medium. This framework provides a physically grounded approximation of long-range electron transfer in biological systems and has been extensively validated for multi-centre redox chains in respiratory proteins. A tunnelling attenuation factor of

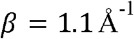

was adopted, consistent with prior experimental and theoretical analyses of protein-mediated electron transfer ^42 18 40^. For each species, inter-centre distances were computed between successive nodes along the canonical electron-transfer pathway from FMN to N2 (FMN → N3 → N1b → N4 → N5 → N6a → N6b → N2), using centroid coordinates defined for each redox cofactor. Distances were calculated as Euclidean separations between centroids:

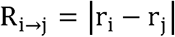

To quantify structural effects on transfer efficiency across species, relative rate perturbations were expressed in logarithmic form:

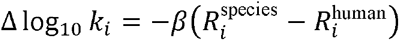

where 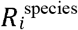 and 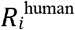denote the distances between equivalent donor–acceptor pairs in the analysed species and in Homo sapiens (5XTD), respectively ^3^. This formulation directly links structural deviations to changes in tunnelling efficiency while remaining independent of absolute prefactor assumptions. The resulting metric enables systematic comparison of electron-transfer topology across evolutionarily diverse Complex I architectures.

### Protein-mediated pathway model and coupling estimation

Electron transfer between redox centers in mitochondrial complex I was further analyzed using a protein-mediated pathway framework that extends beyond direct distance-based tunneling models. While conventional formulations (e.g., Moser–Dutton) approximate electronic coupling as an exponential function of donor–acceptor separation, this approach does not explicitly account for the structural mediation of the protein environment.

To incorporate the effect of the intervening protein matrix, we defined a pathway-dependent tunneling penalty P, which captures the cumulative attenuation of electronic coupling along the protein-mediated electron-transfer route. For a given donor–acceptor pair (N5 → N6a), the pathway penalty was approximated as a structure-derived scalar proportional to the effective tunneling distance through the protein medium:

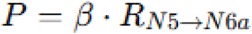

where R_N5_ → N6a is the centroid–centroid distance between the N5 and N6a Fe–S clusters obtained directly from experimentally resolved structures, and β is the effective tunneling decay constant (set to 1.1 Å^−1^, consistent with protein-mediated electron transfer literature). The pathway-derived electronic coupling proxy J_path_ was then defined as:

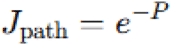

which reflects the attenuation of electronic communication mediated by the protein scaffold. This formulation preserves the exponential distance dependence of tunneling while explicitly framing coupling as a consequence of protein-mediated pathways rather than direct vacuum-like transfer. To relate coupling to functional transport behavior, a pathway-derived rate proxy was estimated assuming a quadratic dependence:

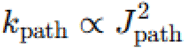

This relation is consistent with non-adiabatic electron transfer theory, in which transfer rates scale with the square of the electronic coupling matrix element. All quantities (P, J_path_, and k_path_) were computed for each species using identical structural definitions, enabling direct cross-species comparison. Statistical relationships between pathway penalty, coupling, and structural distance were evaluated using Pearson and Spearman correlation analyses. Importantly, this pathway-based formulation does not introduce additional empirical fitting parameters beyond the tunneling decay constant, and instead provides a physically grounded, structure-driven approximation of electronic coupling that bridges purely geometric models and full quantum chemical calculations.

### Quantum Hamiltonian and physical interpretation

The redox-chain network was mapped onto a tight-binding Hamiltonian:

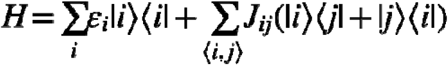

Here, ∣i⟩ represents a localized electronic state at redox center i, and J_ij_ defines quantum coupling between neighboring sites. Site energies were assumed uniform: ϵ_i_ = ϵ_0_, to isolate the effect of coupling heterogeneity. In the present model, site energies were assumed to be uniform in order to isolate the role of structural geometry in governing electron transport. Although differences in redox potentials and reorganization energies are known to influence individual electron-transfer steps, incorporating these factors would introduce additional energetic heterogeneity that could obscure the specific contribution of spatial organization. Our analysis therefore focuses on determining whether geometric constraints alone are sufficient to generate system-level transport limitations. The emergence of a robust structure-dependent bottleneck indicates that spatial organization itself constitutes a major determinant of transport efficiency, while energetic factors are more likely to modulate the magnitude and redistribution of transport flux. This framework reflects the central aim of the present study: to probe structural control of transport rather than energetic fine-tuning.

The uniform-energy model should therefore be interpreted as a deliberately conservative null model. Because geometry-dependent bottlenecks emerge even in the absence of energetic heterogeneity, these transport constraints necessarily originate from spatial organization alone. The inclusion of realistic redox-potential differences would more likely reinforce or redistribute, rather than eliminate, the structurally defined bottleneck. Physically, this Hamiltonian describes a quantum particle propagating through a network with spatially varying tunneling amplitudes, such that structural geometry directly governs the available transport pathways.

### Continuous-time quantum walk dynamics

The coherent evolution of the system is described by:

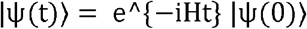

with initial condition:

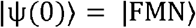

This formulation captures wave-like propagation of the electron across the redox network, allowing interference effects and non-classical transport pathways. Unlike classical hopping models, this approach allows transient delocalization of the electron over multiple sites, which can enhance or suppress transport depending on network structure.

### Identification of structural bottlenecks

The dominant bottleneck was defined as the redox transition exhibiting the minimum electronic coupling: ^37^

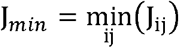

Equivalently, this corresponds to the maximum value of −log_10_(J_ij_ norm). Bottleneck identity and strength were compared across species and distance metrics to assess robustness. ^18,36^

### Open quantum system simulations

Electron transport dynamics were modeled using a reduced open quantum system framework incorporating environmental decoherence. The system was represented as a one-dimensional chain of redox sites with nearest-neighbor coupling.

Transport dynamics were simulated using a Lindblad-type decoherence model:

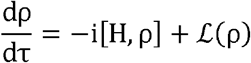

where ρ is the reduced density matrix, H is the tight-binding Hamiltonian constructed from distance-dependent couplings J_ij_, and L(ρ) represents local dephasing processes.^43^

Decoherence was introduced phenomenologically via site-local operators:

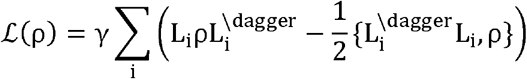

where γ is the dimensionless decoherence strength and Li=∣i⟩⟨i∣. All simulations were performed in dimensionless units, and results are reported in terms of normalized population dynamics.^44^

### Open-system dynamics and Lindblad decoherence

To incorporate environmental effects, the system was treated as an open quantum system using the Lindblad master equation:

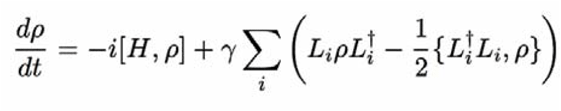

Here, γ represents the dephasing rate and:

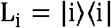

are local dephasing operators. This formulation captures loss of phase coherence without population decay, modeling interactions with a fluctuating protein environment. Physically, increasing γ drives the system from a coherent regime toward a classical hopping limit, enabling exploration of environment-assisted quantum transport.

### Transport observables and efficiency metrics

Transport efficiency was quantified by the population at the terminal site (N2):

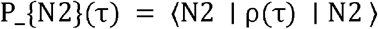

Decoherence sensitivity was defined as:

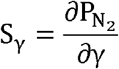

evaluated numerically across a range of decoherence strengths.

### Absorbing sink and irreversible transport

Electron transfer to the quinone site was modeled as an irreversible process using a non-Hermitian sink:

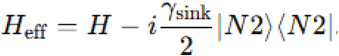

This term removes probability amplitude at the terminal site, representing successful electron transfer. The absorbed population is given by:

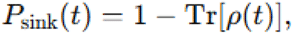

which directly measures transport completion.

### Transport observables and efficiency metrics

Transport efficiency was quantified using:

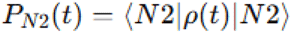

and the characteristic time:

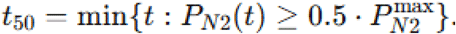

This metric provides a robust measure of transport dynamics that is insensitive to absolute normalization. Additionally, quantum coherence was monitored via:

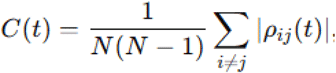

allowing direct assessment of coherence–transport relationships.

### Identification of structural bottlenecks

The dominant bottleneck was defined as:

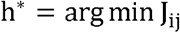

corresponding to the largest tunneling barrier. This definition links structural geometry directly to quantum transport limitation, providing a physically interpretable measure of transport control.

### Numerical implementation and stability considerations

The Lindblad equation was solved using a fourth-order Runge–Kutta integration scheme. Numerical stability was ensured by:

- enforcing Hermiticity:

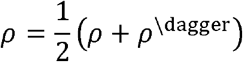

- maintaining trace normalization:

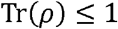

Time evolution was computed over a dense grid to capture multi-scale dynamics.

### Simulation parameters and normalization

All simulations were conducted using dimensionless time τ and decoherence strength γ, enabling comparison of relative transport behavior independent of absolute timescales. Electronic couplings were derived from structure-based distances and scaled such that the strongest coupling within each system was normalized to unity.

### Quantum-walk model of electron transport

Electron transport was modeled as a continuous-time quantum walk on a structure-derived redox network. Redox centers (FMN and Fe–S clusters) were represented as nodes, and electronic couplings between adjacent centers were assigned based on inter-site distances using an exponential decay form:

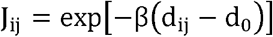

where d_ij_ is the inter-center distance, β is the decay parameter, and d_0_ is a reference distance. The resulting Hamiltonian was constructed as a tight-binding model with nearest-neighbor couplings. The system dynamics were described using the density matrix formalism:

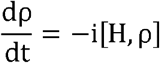

with initial population localized at the FMN site. Time evolution was numerically integrated to obtain site populations and edge-resolved quantum flux. Flux along each edge was computed as:

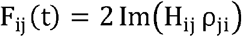

and time-averaged to quantify pathway contributions. This implementation follows a reproducible pipeline for structure-based quantum-walk simulations.

### Decoherence modeling using Lindblad formalism

Environmental effects were incorporated using a Lindblad master equation within a Haken– Strobl dephasing framework:

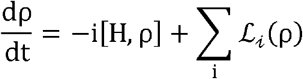

where the Lindblad operators Li=γ∣ i⟩⟩ i∣ introduce pure dephasing with rate γ. An irreversible sink was introduced at the terminal N2 site to model electron trapping: ^45^

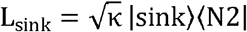

The Liouvillian superoperator was constructed using standard vectorization:

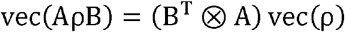

and the system was propagated using eigendecomposition of the Liouvillian. For each species, decoherence strength γ was varied over several orders of magnitude (10^−4^–10^2^), and the population transferred to the sink at a fixed observation time was computed.

Decoherence sensitivity was quantified as:

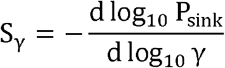

evaluated near γ≈1, allowing comparison across systems. All simulations were performed using a 9-state model (8 redox sites + sink) with experimentally derived coupling strengths.

### Residue-mediated alternative pathway analysis

To identify alternative electron-transfer pathways, a local network was constructed around the N5–N6a interface using atomistic structural data. Residues located within a geometric corridor between the two redox centers were selected based on minimum atom–atom distances:

- d_N5–res_ + d_res–N6a_<d _cutoff_
- and proximity constraints to either redox site

Selected residues were treated as intermediate nodes, forming a network that includes both direct (N5–N6a) and residue-mediated connections. Couplings between nodes were assigned as:

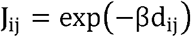

with separate treatment for direct redox coupling and bridge-mediated interactions. Quantum transport on this extended network was simulated using a coherent continuous-time quantum walk. Bridge residues were assigned positive on-site energy offsets to represent virtual tunneling intermediates rather than trapping states. To quantify pathway competition, the direct N5–N6a coupling was systematically scaled, and the resulting redistribution of quantum flux was evaluated. The relative contribution of alternative pathways was defined as:

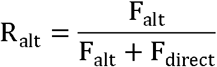

where F_alt_ and F_direct_ denote time-averaged flux through alternative and direct pathways, respectively. Flux values were obtained by integrating the absolute quantum flux over time for each edge. This approach enables identification of dominant transport routes and quantification of flux redistribution under varying bottleneck strengths

### Visualization

All figures were generated using Python (NumPy, SciPy, Matplotlib):

### Scope and limitations

The present model isolates the role of structure and quantum dynamics under several assumptions, including uniform site energies and nearest-neighbor coupling. While these simplifications neglect detailed electronic structure and long-range interactions, they allow identification of dominant physical mechanisms.

Importantly, the model should be interpreted as a minimal quantum transport framework that captures the interplay between geometry, coherence, and dissipation, rather than a full ab initio simulation.

## Declarations

### Ethics approval and consent to participate

N/A

### Consent for publication

N/A

### Availability of data and materials

The analysis codes and datasets used and/or generated during the current study are available from Dr. Ji-Yong Sung upon reasonable request. Interested researchers may contact Dr. Sung via email at to obtain access to the relevant materials.

### AI Use Declaration

AI tools were used only for English grammar correction and language polishing.

### Authors’ contributions

Conceptualization & Investigation: JYS, JHC; Methodology: JYS; Data analysis: JYS; Writing-original draft: JYS; Writing-review & editing: JYS, JHC; Supervision: JYS, JHC; Project administration: JYS, JHC; Funding acquisition: JHC; Interpretation of the results; JYS, JHC. All authors have read and agreed to the published version of the manuscript.

## Acknowledgements

N/A

---

**Supplementary Fig. S1.**
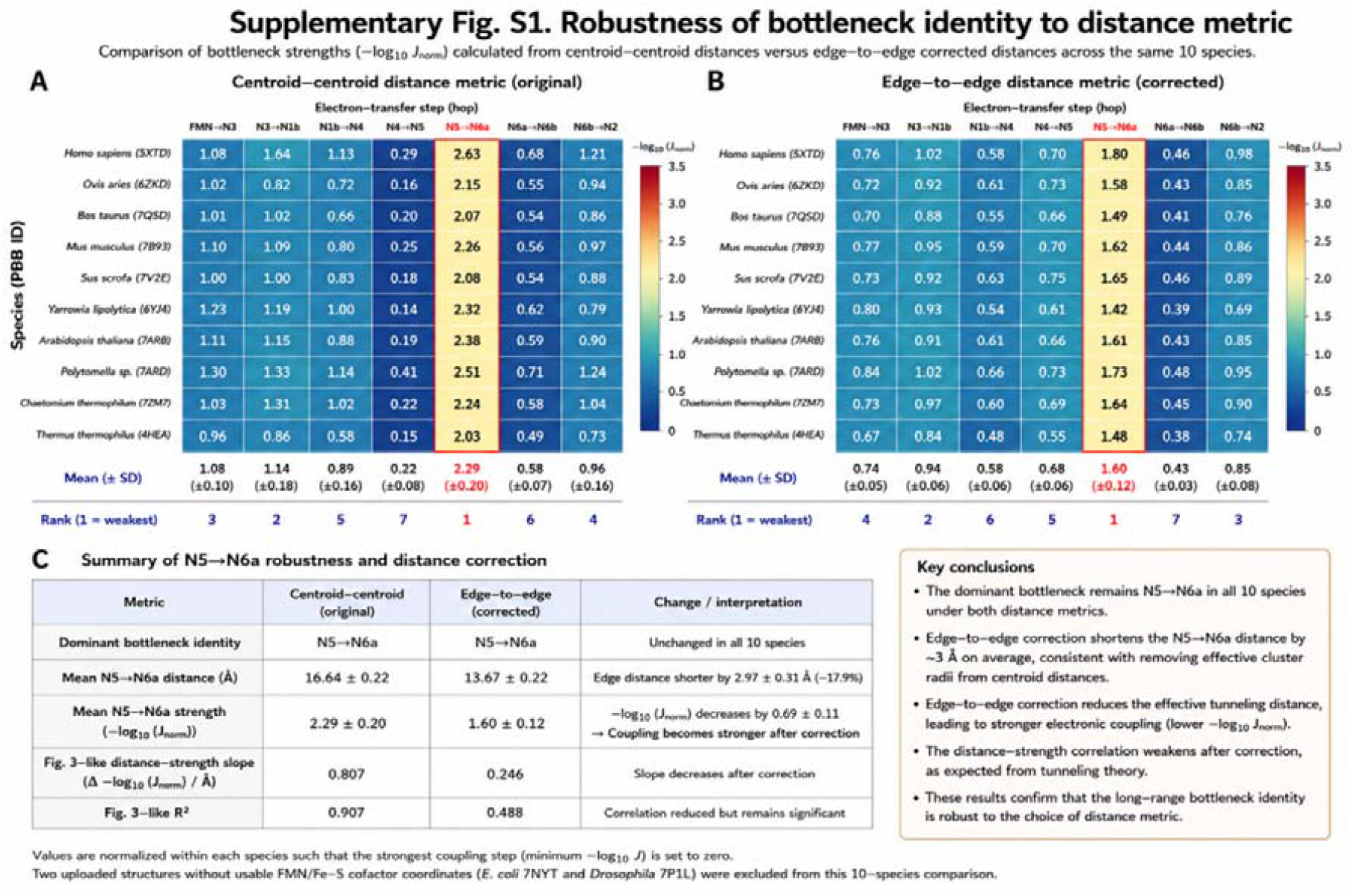
Robustness of bottleneck identity to distance metric. (A) Heatmap of bottleneck strengths (−log_10_J_norm) calculated using centroid–centroid distances for electron-transfer (ET) steps across 10 Complex I structures from different species. ET couplings were normalized within each species, such that the strongest coupling step was assigned a value of 0. The N5–N6a step consistently exhibited the highest bottleneck strength across all species, indicating the weakest effective electronic coupling in the ET chain. Mean ± SD values across species and rank ordering of bottleneck strengths are shown below the heatmap. (B) Heatmap of bottleneck strengths recalculated using edge-to-edge corrected distances, which account for van der Waals radii and reduce the effective tunneling distance between cofactors. Although absolute bottleneck strengths decreased after correction, the N5–N6a step remained the dominant bottleneck in all analyzed species. Mean ± SD values and rank ordering are shown below the heatmap. (C) Summary comparison between centroid–centroid and edge-to-edge distance metrics. Edge-to-edge correction shortened the average N5–N6a distance from 16.64 ± 0.22 Å to 13.67 ± 0.22 Å and reduced the normalized bottleneck strength from 2.29 ± 0.20 to 1.60 ± 0.12. Consistent with the exponential distance dependence of electronic coupling, correction of the effective tunneling distance weakened the distance–strength correlation slope but preserved the overall bottleneck identity. These results demonstrate that the long-range bottleneck assignment in Complex I ET pathways is robust to the choice of distance metric. Two structures lacking complete FMN/Fe–S cofactor coordinates (Escherichia coli 7NYT and Drosophila melanogaster 7P1L) were excluded from this analysis.

**Supplementary Fig. S2.**
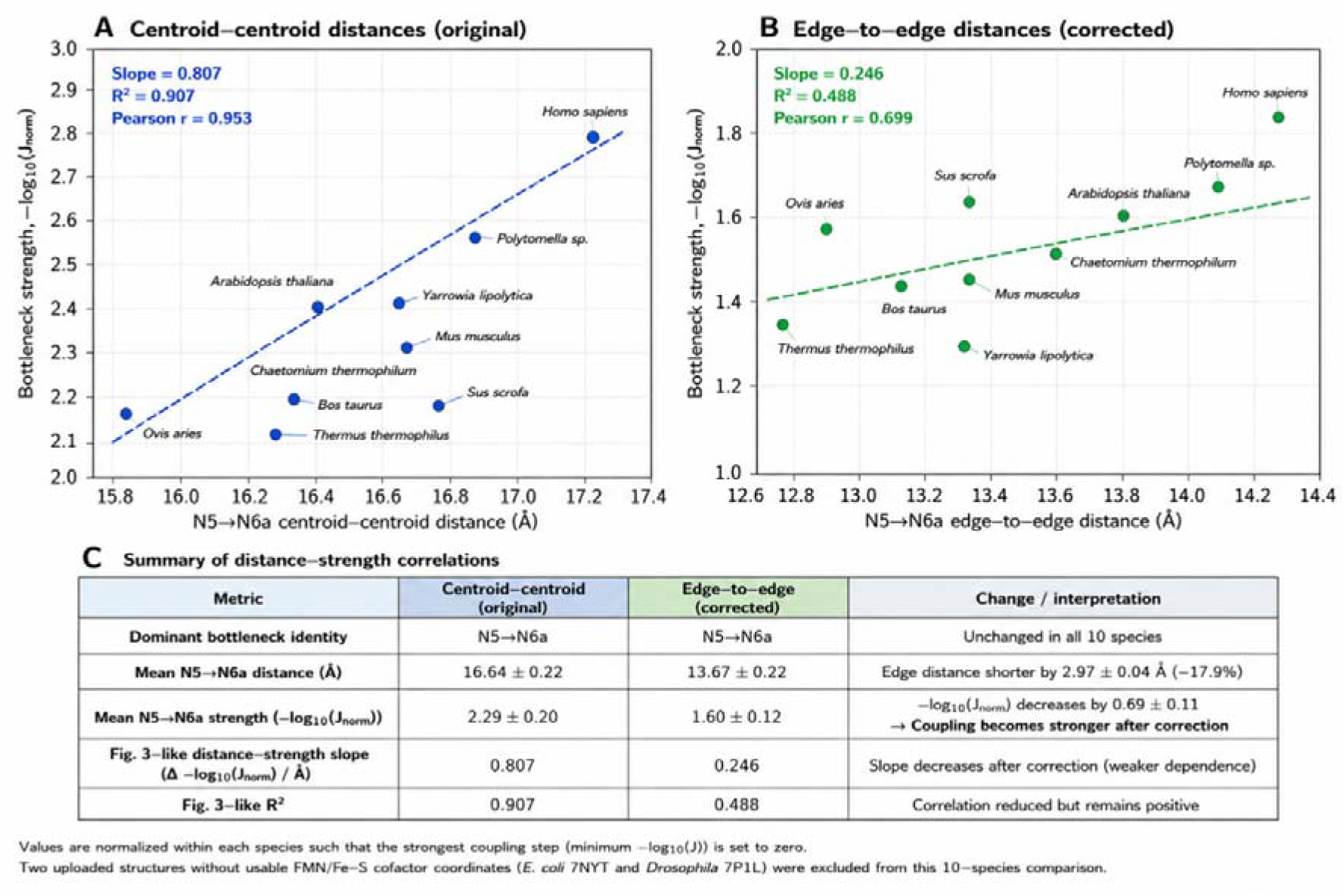
Effect of edge-to-edge distance correction on distance–strength correlations in Complex I electron transfer pathways. (A) Correlation between the N5– N6a bottleneck distance and normalized bottleneck strength (−log_10_J_norm_) calculated using centroid–centroid distances across 10 species. A strong positive linear relationship was observed (slope = 0.807, R^2^ = 0.907, Pearson r = 0.953), consistent with the exponential distance dependence of electronic coupling predicted by electron-tunneling theory. Species labels indicate the corresponding Complex I structures used in the analysis. (B) Correlation between N5–N6a bottleneck distance and normalized bottleneck strength after applying edge-to-edge distance correction. Corrected distances account for van der Waals radii and reduce the effective tunneling separation between cofactors. Although the overall distance dependence became weaker after correction (slope = 0.246, R^2^ = 0.488, Pearson r = 0.699), the correlation remained positive, indicating that ET bottleneck behavior is still governed primarily by inter-cofactor separation. (C) Summary table comparing centroid–centroid and edge-to-edge distance metrics. Edge-to-edge correction shortened the average N5–N6a distance from 16.64 ± 0.22 Å to 13.67 ± 0.22 Å (−17.9%) and reduced the normalized bottleneck strength from 2.29 ± 0.20 to 1.60 ± 0.12. The decrease in slope after correction indicates weaker apparent distance dependence due to reduced effective tunneling lengths, while the dominant bottleneck identity (N5–N6a) remained unchanged across all analyzed species. Values were normalized within each species such that the strongest coupling step (minimum −log_10_J) was set to zero. Two uploaded structures lacking complete FMN/Fe–S cofactor coordinates (Escherichia coli 7NYT and Drosophila melanogaster 7P1L) were excluded from this comparison.

**Supplementary Fig. S3.**
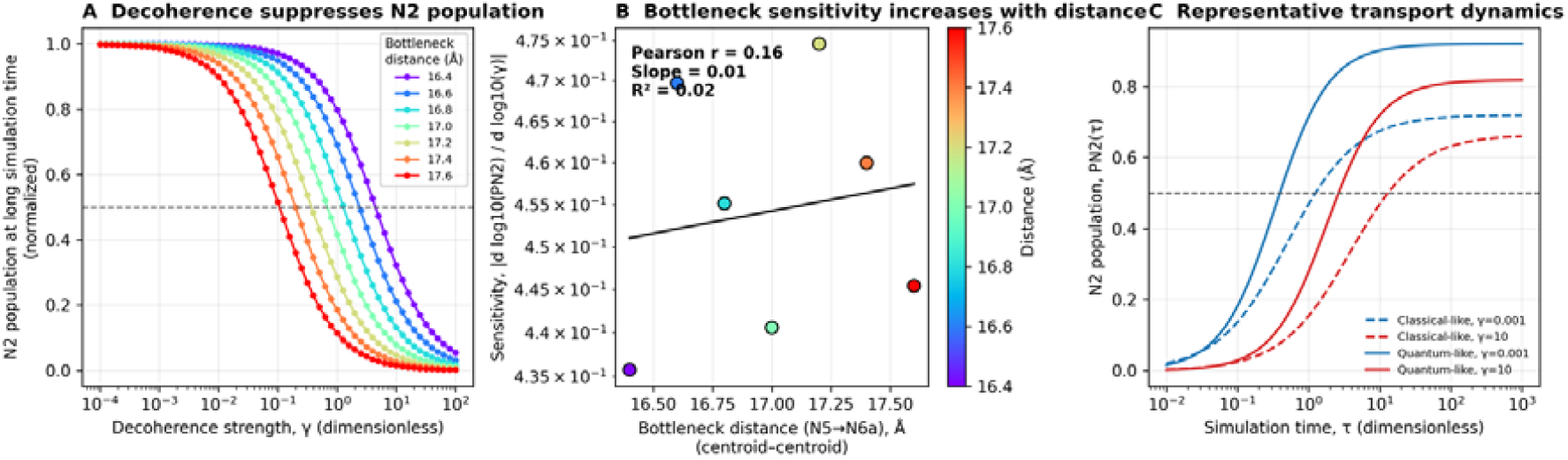
Decoherence-dependent suppression of N2 population and bottleneck sensitivity in Complex I electron transport. (A) Final N2 population as a function of decoherence strength (γ) for different N5–N6a bottleneck distances. Increasing decoherence progressively suppresses long-time N2 population, with larger bottleneck distances exhibiting earlier and stronger population collapse. This behavior indicates that weak electronic coupling at long tunneling distances enhances sensitivity to environmental decoherence. (B) Relationship between bottleneck distance (N5–N6a separation) and decoherence sensitivity, quantified as the absolute change in N2 population (|Δlog_10_PN2|) under increasing γ. A weak positive trend was observed (Pearson r = 0.16, slope = 0.01, R^2^ = 0.02), suggesting that longer tunneling distances modestly increase transport vulnerability to decoherence, although substantial species-specific variability remains. Point colors represent centroid–centroid bottleneck distances. (C) Representative time evolution of N2 population under classical-like (γ = 100) and quantum-like (γ = 0.001) transport regimes. Quantum-like transport exhibits faster initial population transfer and earlier saturation, whereas strong decoherence slows transport kinetics and reduces steady-state accumulation. Blue and red curves represent shorter and longer bottleneck distances, respectively. The dashed horizontal line indicates PN2 = 0.5. Simulation time (τ) and decoherence strength (γ) are shown in dimensionless units.

