## Supplementary figures and images for "Quantum transport in mitochondrial complex I is governed by a conserved structural bottleneck"

### Supplementary Fig3

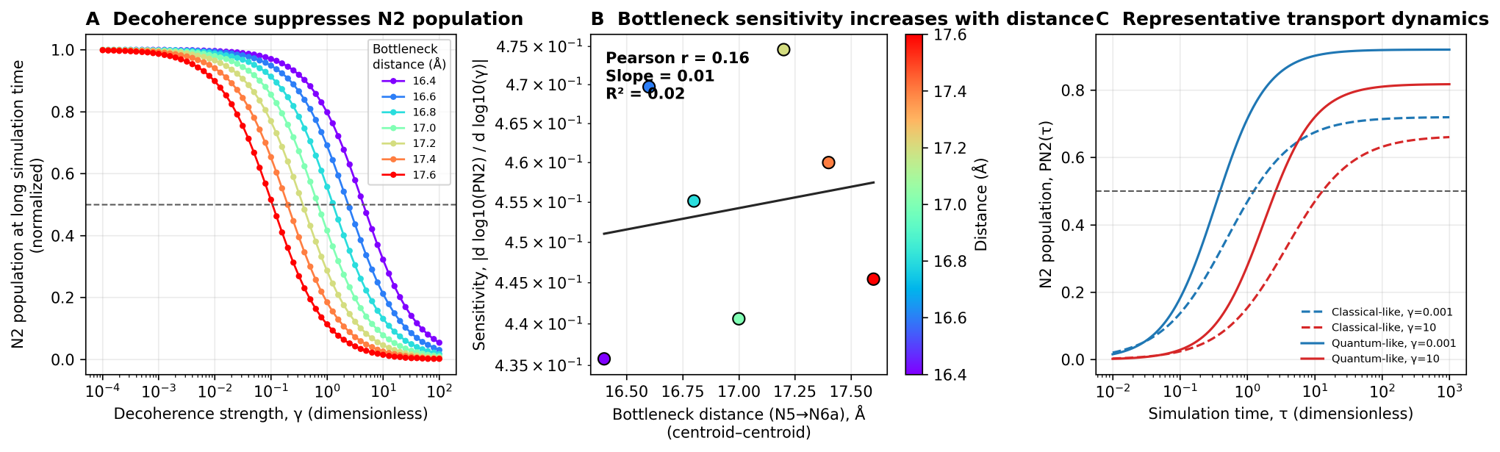

### Supplementary Fig. 1

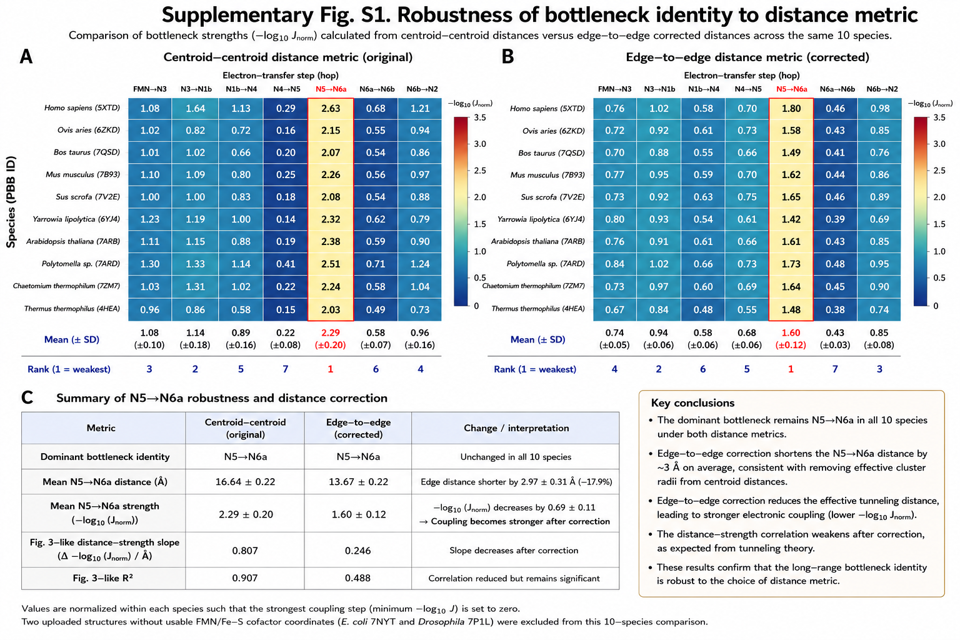

### Supplementary Fig. 2

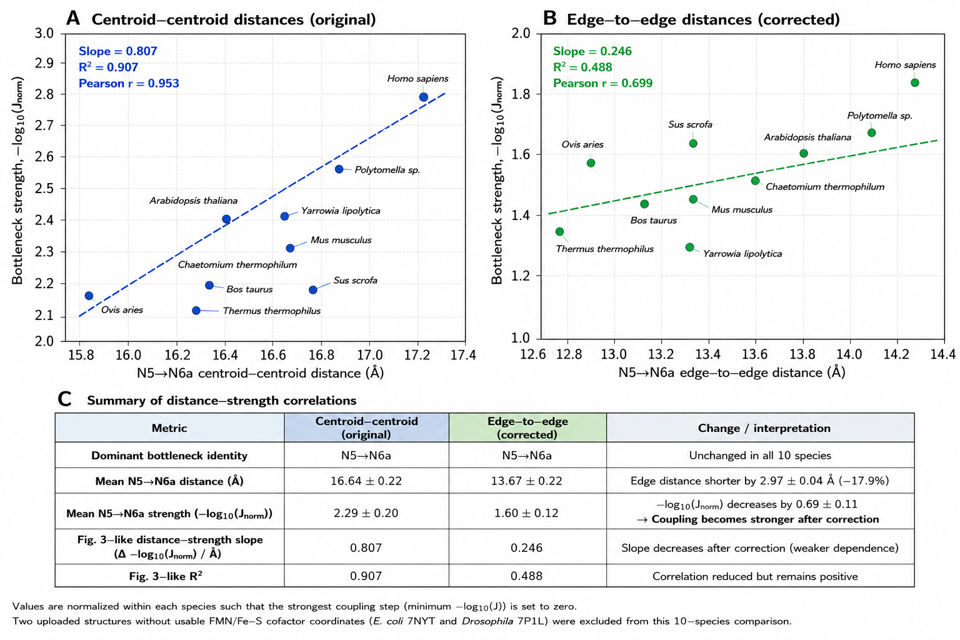
